# Coupling water fluxes with cell wall mechanics in a multicellular model of plant development

**DOI:** 10.1101/511717

**Authors:** Ibrahim Cheddadi, Michel Génard, Nadia Bertin, Christophe Godin

## Abstract

The growth of plant organs is a complex process powered by osmosis that attracts water inside the cells; this influx induces simultaneously an elastic extension of the walls and pressure in the cells, called turgor pressure; above a threshold, the walls yield and the cells grow. Based on Lockhart’s seminal work, various models of plant morphogenesis have been proposed, either for single cells, or focusing on the wall mechanical properties. However, the synergistic coupling of fluxes and wall mechanics has not yet been fully addressed in a multicellular model. This work lays the foundations of such a model, by simplifying as much as possible each process and putting emphasis on the coupling itself. Its emergent properties are rich and can help to understand plant morphogenesis. In particular, we show that the model can display a new type of lateral inhibitory mechanism that could contribute to the amplification of growth heterogeneities, essential for shape differentiation.

**Significance Statement:** Plant cells are surrounded by a rigid wall that prevents cell displacements and rearrangements as in animal tissues. Therefore, plant morphogenesis relies only on cell divisions, shape changes, and local modulation of growth rate. It has long been recognized that cell growth relies on the competition between osmosis that tends to attract water into the cells and wall mechanics that resists to it, but this interplay has never been fully explored in a multicellular model. The goal of this work is to analyze the theoretical consequences of this coupling. We show that the emergent behavior is rich and complex: among other findings, pressure and growth rate heterogeneities are predicted without any ad-hoc assumption; furthermore the model can display a new type of lateral inhibition based on fluxes that could complement and strengthen the efficiency of already known mechanisms.

---

Plants grow throughout their lifetime at the level of small regions containing undifferentiated cells, the meristems,located at the extremities of their axes. Growth is poweredby osmosis that tends to attract water inside the cells. The corresponding increase in volume leads to simultaneous tension in the walls and hydrostatic pressure (so-called turgor pressure)in the cells. Continuous growth occurs thanks to the yielding of the walls to these stretching forces [1–3].

This interplay between growth, water fluxes, wall stress and turgor was first modelled by Lockhart in 1965[4], in the context of a single elongating cell. Recent models focused on how genes regulate growth at more integrated levels [5–9]. To accompany genetic, molecular, and biophysical analyses of growing tissues, various extensions of Lockhart’s model to multicellular tissues have been developed. The resulting models are intrinsically complex as they represent collections from tens to thousands of cells in 2- or 3-dimensions interacting with each other. To cut down the complexity, several approaches abstract organ multicellular structures as polygonal networks of 1D visco-elastic springs either in 2D [7,10–12] or in 3D [6, 13] submitted to a steady turgor pressure. Other approaches try to represent more realistically the structure of the plant walls by 2D deformable wall elements able to respond locally to turgor pressure by anisotropic growth [8, 14, 15].

Most of these approaches consider turgor as a constant driving force for growth, explicitely or implicitly assuming that fluxes occur much faster than wall synthesis. Cells then regulate the tissue deformations by locally modulating the material structure of their walls (stiffness and anisotropy) [6, 16–20]. However, the situation in real plants is more complex: turgor heterogeneity has been observed at cellular level [21, 22], which challenges the assumption of very fast fluxes. As a matter of fact, the relative importance of fluxes or wall mechanics as limiting factors to growth has fuelled a long standing debate [3, 23] and is still an open question. Moreover, from a physical point of view, pressure is a dynamic quantity that permanently adjusts to both mechanical and hydraulic constraints, which implies that a consistent representation of turgor requires to model both wall mechanics and hydraulic fluxes.

The aim of this article is to explore the potential effect of; coupling mechanical and hydraulic processes on the properties of the “living material” that corresponds to multicellular; populations of plant cells. To this end, we build a model; that describes in a simple manner wall mechanics and cell structure, but do not compromise on the inherent complexity of considering a collection of deformable object hydraulically and mechanically connected.

The article is organized as follows (see Fig. 1): we first recall the Lockhart-Ortega model and its main properties. Then we explore two simple extensions of this model: first we relax the constraint of uniaxial growth in the case of a single polygonal cell; then we study how two cells hydraulically connected interact with each other. Finally we describe our multicellular and multidimensional model and numerically explore its properties.

**Fig. 1.**
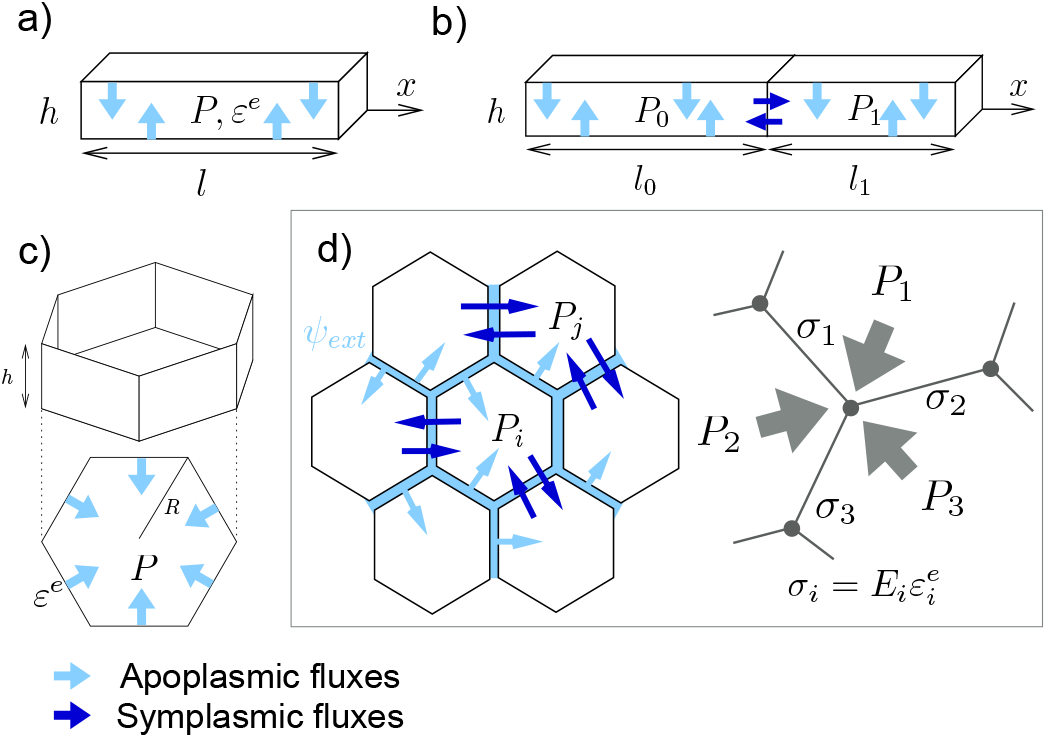
Hierarchy of models presented in this article. Main variables are turgor *P* and elastic deformation *ε^e^*. **a)** Lockhart-Ortega model: uniaxial growth in the *x* direction of a cylindrical cell of length l; the section perpendicular to *x* is a square of side *h*. **b)** two cells extension, both growing along *x*; **c)** 2D extension of a single cell growth; **d)** Multicellular, multidimensional model; left: fluxes, right: mechanical equilibrium; the stress *σ* is proportionnal to the elastic deformation *ε^e^*; *E* is the elastic modulus.

## The Lockhart model

In 1965, Lockhart [4] derived the elongation of a cylindrical plant cell by coupling osmosis-based fluxes and visco-plastic wall mechanics. Ortega [24] extended this seminal model to include the elastics properties of the cell walls. We recall here the main properties of this model, see Fig. 1a for the geometrical configuration.

### Cell wall elongation

It is expressed as a rheological law [4, 24]: the total strain rate of the walls 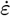 is decomposed into the sum of a plastic and an elastic strain rate:

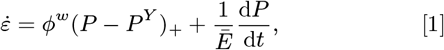

where the extensibility *ϕ^w^* (inverse of a viscosity) describes the ability of the cell to synthesize wall material, and *Ē* is an effective elastic modulus. Here, *ϕ^w^* and *Ē* both depend on cell wall thickness. The notation (*x*) + denotes *x* if *x* > 0 and 0 otherwise for any real number *x*.

### Water uptake

Lockhart described water uptake by the cell as a flux through a semi-permeable membrane characterized by its surface *A* and its permeability *L^a^*. Assuming the membrane is perfectly impermeable to solutes, the rate of volume change is the result of a difference between the water potential Ψ of the cell and Ψ_*ext*_ of its exterior [25]:

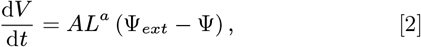

The cell water potential Ψ = *P − π* results from the antagonistic effect of the cell hydrostatic pressure *P* that tends to expel water from the cell and its osmotic pressure *π* that tends to attract water inside the cell. In the case of a single solute of concentration c, we have *π* = *RTc* where *R* is the ideal gas constant and *T* the temperature. Let us denote 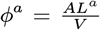 which has the same dimension as *ϕ^w^*. Assuming that the fluxes occur mostly on the lateral surface, the ratio *A/V* is constant in the configuration of a cylindrical cell. After division by *V*, Eq. (2) turns into:

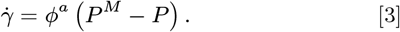

where *P^M^* = Ψ*_ext_*+*π* quantifies the power of the osmotic pump: it is positive if *π* is high enough to overcome the negative water potential of the exterior of the cell. Growth 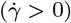 implies *P* < *P^M^* and hence *P^M^* is an upper bound for turgor, above which the cell would lose water to the exterior. The additional condition for growth *P* > *P^Y^* (see above) requires *P^M^* > *P^Y^*: growth is possible only when the osmotic pump is able to overcome the mechanical resistance of the walls.

In order to keep the analysis as simple as possible, we take here and in the remaining of the article *P^M^* constant with time and homogeneous among the cells, which corresponds for instance to constant *π* and Ψ*_ext_*. This choice will be commented in the discussion section.

### Coupling hydraulics and mechanics for a single cell

Equating the expressions of strain rate *ε* from Eq. (1) and relative growth rate 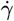 from Eq. (3) ensures that the requirements for water uptake and yield of the cell wall are simultaneously satisfied. This means that turgor *P*, that is present in both equations, has to be adjusted to satisfy both hydraulic and mechanical constraints. The resolution of the model is detailed in Supplementary Information (SI), Eqs. (S3)-(S4). The time dependent solutions can be analytically determined and we find that *P* and 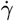 converge towards a stationary solution 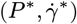: first, *P\** writes

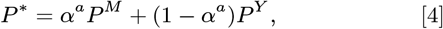

where

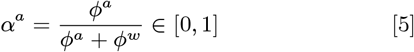

measures the relative importance of *ϕ^a^* compared to *ϕ^w^*. In the limit *ϕ^a^* ≪ *ϕ^w^* (*α^a^* = 0), any excess of turgor above the threshold is relaxed by cell wall synthesis and turgor is minimal at *P* = *P^Y^*. Conversely, in the limit *ϕ^w^* ≪ *ϕ^a^* (*α^a^* = 1), the wall synthesis is not able to relax turgor, which reaches then its maximal value *P* = *P^M^*. Second, the expression of the relative growth rate is:

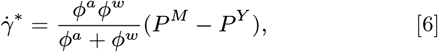

or equivalently: 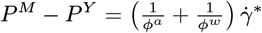. This equation is the analog of Ohm’s law Δ*U* = (*R*_1_ + *R*_2_)*I* with two resistors *R*_1_ = 1/*ϕ^a^* and *R*_2_ = 1/*ϕ^w^* in series: growth can be limited by either hydraulic conductivity or wall synthesis.

Link with wall rheology. Wall expansion law (Eq. (1)) can be equivalently described as a function of wall stress *σ* rather than cell turgor *P*: in the cylindrical geometry of the Lockhart-Ortega model, we find (see SI for the calculations) 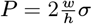, where *w* is the width of the walls and *h* their height. Thanks to this relation, Eq. (1) translates into 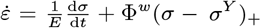, where 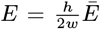 (resp. 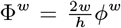) is the intrinsic elastic modulus (resp. extensibility) of the walls. Let *ε^e^ = σ/E* be the so-called elastic deformation of the walls. It is dimensionless and can be measured from the image analysis of experiments, without the knowledge of the elastic modulus. The wall rheology is then described as follows:

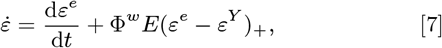

where *ε^Y^* = *σ^Y^/E* is the threshold elastic deformation. Note that 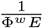 can be interpreted as the characteristic time of wall synthesis.

## Multidimensional and multicellular models

A multicellular extension of the Lockhart-Ortega model adapted to the study of morphogenesis requires first to relax the constraint of uniaxial growth and allow multidimensional geometries, and second is complexified by the possibility of fluxes between cells. We study separately the effect of each of these extensions before presenting the complete model.

### First extension: Multidimensional growth

In order to keep the analysis as simple as possible, we study here the expansion of a single 2D cell whose shape is a regular polygon with *n* edges (see Fig. 1c). This model allows to evaluate the effect of a varying surface/volume ratio compared to the Lockhart-Ortega model where this ratio is constant. The fluxes are described in the same way as for Lockhart’s model (Eq. (2)) but wall synthesis is described with Eq. (7), as a function of elastic deformation instead of turgor. We find (see SI for detailed calculations) that the relation between cell turgor and wall stress becomes 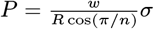 where *R* is the cell radius. In contrast with the Lockhart-Ortega model, the ratio *P/σ* is no more constant as cell grows, and the turgor vanishes at long times if the stress remains in the order of magnitude of the threshold. Note also that for a given stress the turgor decreases with the number of edges n. Therefore, the yield turgor *P^Y^*depends both on *n* and *R* and is not a well defined parameter. It suggests also that cells with less neighbours should have a higher turgor, as experimentally observed in [21, 22].

The prediction of growth rate requires a numerical resolution of the model (see SI). The parameters are chosen to ensure a turgor of the order of 0.5 MPa and a relative growth rate of the order of 2% per hour, using the predictions Eq. (4) and Eq. (6). First let’s examine the case of a cell of initial radius *R* = 10*μm* for which wall synthesis is the limiting factor to growth (case *α^a^* = 0.9 in SI, fig. S2). We find that it results initially in an accelerating growth (the bigger the cell, the faster the growth), much faster than predicted by the Lockhart model, during which the elastic deformation of the walls can reach values up to 20%. The ratio area/surface = 1/*R* decreases with growth and there is less and less water available compared to the volume; as a consequence, the relative growth rate vanishes at long times after this initial accelerating phase.

In the case where the fluxes are already limiting in the initial state (case *α^a^* = 0.1 in SI, Fig. S2), the initial behaviour is closer to the predictions of the Lockhart model but the relative growth rate still vanishes at long times.

Altogether, these results show that a non constant surface/volume ratio deeply modifies the behavior of the model compared to the Lockhart model. In particular, flux and wall synthesis as limiting factors fro growth are no more equivalent.

### Second extension: Multicellular growth

Then, we study a simple multicellular extension of the Lockhart-Ortega model where two cylindrical cells *i* = 0, 1 are in contact through one of their wall (see Fig. 1b). The cells can absorb water from their lateral surface and in the meantime exchange water with each other through their common wall. We look for stationary solutions: 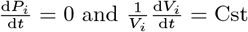.

We set for both cells a common value of *P^M^*, *L^a^* and *ϕ^w^*, while the value of the yield turgors 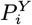 can differ; this corresponds for instance to a heterogeneity of wall elastic modulus or yield deformation. For the sake of convenience, we refer to fluxes between cells as symplasmic fluxes, characterized by a water conductivity *L^s^*, and to fluxes from the water source as apoplasmic fluxes, characterized by a water conductivity *L^a^*. Assuming that the symplasmic fluxes occur through plasmodesmata that are permeable to both water and solutes, the flux equation writes

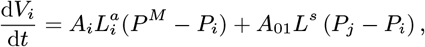

where *j* = 1 − *i*, and *A*_01_ is the surface of the common wall of cells 0 and 1. We introduce the number *ϕ^s^* = 2*A*_01_*L^s^*/*V_i_* which has the same dimension as *ϕ^a^* and *ϕ^w^*. In order to allow an analytical resolution of this set of equations, we assume *ϕ^s^* to be constant with time, and consider it in this section as a parameter of the model. Thus, we have

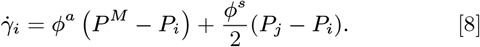

We introduce the dimensionless number

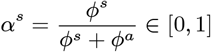

which represents the relative importance of symplastic fluxes with respect to apoplastic ones. We combine this flux equation with the growth equation Eq. (1) and find analytical solutions for any values of the parameters (see SI). We use here the following set of control parameters:

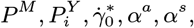

and fix the value 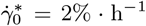; this way, the parameters space to explore is reduced to (*P^M^*, *P^Y^*, *α^a^*, *α^s^*). When *α^s^* = 0, the cells are completely isolated one from another and reach turgors 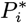 and growth rates 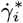 as predicted by the Lockhart model (Eq. (4) and Eq. (6)). In particular, the condition 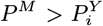 ensures that each cell is growing. When *α^s^* > 0, the fluxes between cells modify this behaviour. We restrict to the case 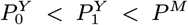, which corresponds to less mechanical constraints on cell 0 than cell 1; therefore we can expect *P*_1_> *P*_0_ and 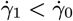. The calculations show a complex non linear behaviour that is illustrated in Fig. 2, in which the parameters subspace (*α^a^*, *α^s^*) is explored for given values of 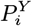 and *P^M^* (detailed calculations are provided in SI). Let 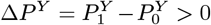 be the difference of the two yield turgors and 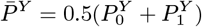 their average; we also introduce the dimensionless number

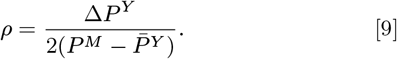

**Fig. 2.**
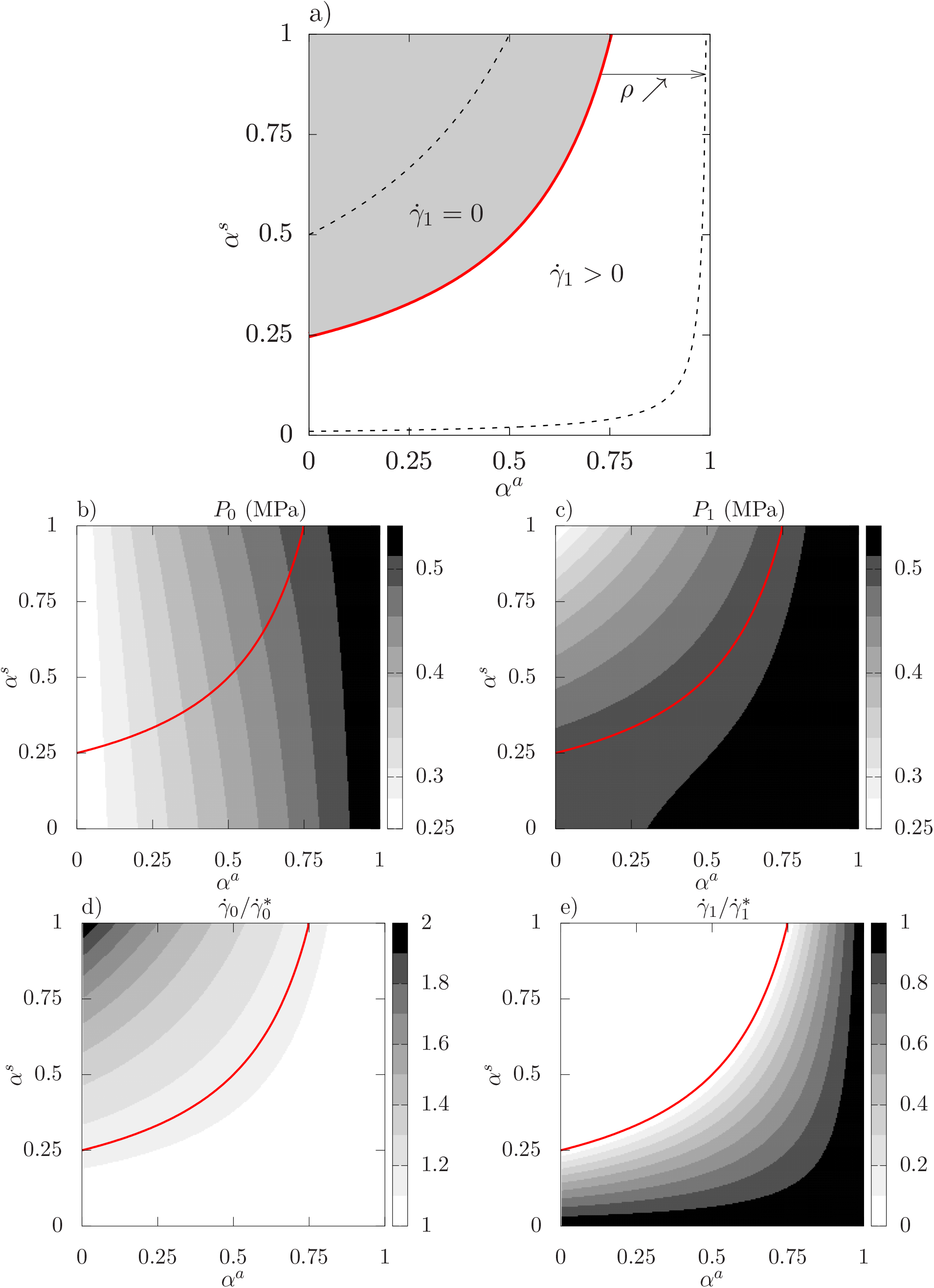
Analytical resolution of the two cells model, properties of the solution in the parameters space *α^a^* × *α^s^*; **a)** delimitation of the two zones 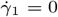 and 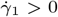: the red thick solid line 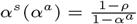 corresponds to *ρ* = 0.75. The two black thin dashed lines correspond to the values *ρ* = 0.5 and 0.99. **b-c)** Turgors *P*_0_ and *P*_1_ for *ρ* = 0.75. **d-e)** relative growth rates 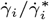 for *ρ* = 0.75.

Note that the hypothesis 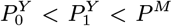 is equivalent to *ρ* ∈]0, 1[.

We find that the subspace (*α^a^*, *α^s^*) can be divided in two main regions separated by the curve 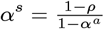 (see Fig. 2a): surprisingly, in the region 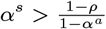, only cell 0 is growing (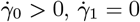, and equivalently 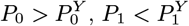). Hence, the growth of cell 1 is inhibited by fluxes with cell 0. Conversely, in the region 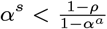 both cells are growing (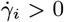 and equivalently 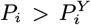). The size of the region 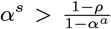 increases with *ρ* and fills the whole square [0, 1] × [0, 1] when *ρ* → 1; such values can be reached when Δ*P^Y^* is large and / or *P^M^* is close to 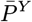.

More quantitatively, Figs. 2d-e) show that 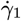 is always below 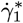, while 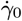 is always above 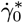 and can reach up to twice this value. Furthermore, maximal values of 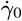 coincide with minimal values of 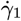: this confirms quantitatively that the growth of the cell with less favorable mechanical condition is slowed down if not inhibited by the growth of its neighbour. This shows also that the growth rate heterogeneity is amplified by fluxes.

Turgor heterogeneity is also affected by fluxes (see Figs. 2b-c): when *α^s^* is close to zero, the cells are hydraulically isolated and their turgors vary with *α^a^* as predicted by Lockhart model (Eq. (4)), this is where the turgor heterogeneity is maximal. Conversely, when *α^s^* is close to 1, there is no hydraulic resistance between the two cells and the two turgors are equal. Between these two limits, *P*_0_ is only slightly affected and remains in the 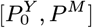 interval; conversely, *P*_1_ is dramatically affected as it shifts from the interval 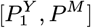 when *α^s^* = 0 to the interval 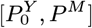 when *α^s^* = 1. Therefore, as 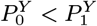, there is a region where 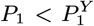 which corresponds to the region 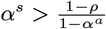, where cell 1 is not growing.

Finally, we have seen that intercellular fluxes tend to increase (resp. decrease) growth rate (resp. turgor) heterogeneities; the cell with less mechanical constraints takes control over the other one and imposes its turgor, which can lead the other one to stop growing. The growing cell then benefits from the water resources of the other cell and its growth is all the more increased.

### Generalization: a multidimensional and multicellular model of growth

We consider (see Fig. 1d) a collection of *N* cells that form a (non necessarily regular) 2D mesh with a fixed topology (distribution of neighbours) as is the case with plant tissues when no division occurs.

The cell walls rheology is described by the visco-elasto-plastic law (Eq. (7)) of the Ortega model and the fluxes toward a cell *i* are described as in the simple multicellular model presented above:

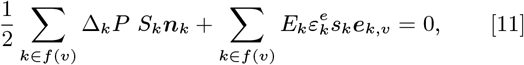

where *n*(*i*) is the set of neighbours of cell *i*, *A_ij_* is the area of the common wall with cell *j*, 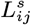 its permeability (it is symmetric: 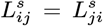), and 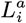 is the permeability of the lateral walls to the supply of water.

The last missing part to obtain a closed set of equation is the mechanical equilibrium, that allows to link cells turgors, walls tensions, and geometry. Contrary to the cases studied above, no explicit expression of turgors as a function of stresses can be obtained and the equilibrium has to be solved at each time step. Let *P_i_* be the turgor pressure in each cell *i*. The tissue being at every moment in a quasi-static equilibrium, pressure forces on wall edges and elastic forces within walls balance exactly at each vertex *v*:

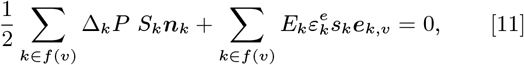

where *f*(*v*) is the set of faces adjacent to junction *v*, Δ*_k_P* = *P*_*k*_1__ − *P*_*k*_2__ is the pressure jump across face *k*, with *k*_1_ < *k*_2_ being indices of the cells across face *k*, *S_k_* = *hl_k_* is the area of the face *k* on which pressure is exerted, *n_k_* is the normal vector to face *k*, oriented from cell *k*_1_ to cell *k*_2_, and *s_k_* = *hw* is the cross-section area of the face, on which the elastic stress is exerted; finally, *e_k,v_* is the unit vector in the direction of face *k*, oriented from junction *v* to the other end of face *k*.

### Coupling mechanical and hydraulic models

In the Lockhart-Ortega model, the compatibility between wall enlargement and cell volume variation is automatically enforced through the geometrical constraint of uni-directional growth that leads to the identity between the relative growth rate of the cell and the strain rate of the walls. In contrast, in the multicellular model, this identity is no longer true. One has to solve the closed set of equations Eq. (7)-Eq. (10)-Eq. (11) with respect to the unknowns *X*, *P*, and *ε^e^*.

Despite its apparent simplicity, the problem to be solved is not straightforward as water fluxes induce potentially long range interactions. In this respect, it differs from most vertex-based models (*e.g* [11, 26]) where turgor is an input of the model. The numerical resolution required the development of an original algorithm (see SI) implemented in an in-house code.

### Numerical experiments: growth of primordia in the shoot apical meristem (SAM)

The properties of this model cannot be as thoroughly studied as those of the simpler models presented above, first because of the numerical cost of the resolution, but above all because it allows an infinite variety of geometries and spatial distribution of its parameters. We present here a numerical experiment that illustrates on the one hand how the properties of the simple multidimensional and multicellular submodels are combined in the generalized model; in turn the study of these models helps us to anticipate the properties of the generalized model. And on the other hand, we show that this model is readily applicable to the study of systems of biological interest.

Growth heterogeneities can be triggered by the local modulation of the mechanical properties of the cell walls [27]. In SAMs, new organs are initiated by a local increase in growth rate that leads to the appearance of small bumps. Measurements show that physico-chemical properties of walls are modified so that mechanical anisotropy and elastic modulus are decreased. In our 2D model, we can explore what effect a local softening of the walls has on growth rate and turgor heterogeneities; based on our previous analysis of the model in simple configurations, we expect that the growth heterogeneities will be maximal for parameters such that the growth is restricted by fluxes rather than wall synthesis (low *α^a^*), cell-cell conductivity is large, and the walls deformations are just above the growth threshold, which can be enforced by a low value of the osmotic pressure (yet large enough to ensure growth). The set of parameters (REF) is chosen according to these criteria; then we explore the effect of a higher *α^a^*((ALPHA+) set) and lower cell-cell conductivity ((CC-) set) that should both decrease the growth heterogeneities, and also test the effect of a lower osmotic pressure ((PM-) set) that should conversely increase the growth heterogeneity. See table 1 in SI for the values of the parameters corresponding to these sets and SI for more precise explanations.

We build a mesh made primarily of hexagons (see Fig. 3a) and first let it grow with homogeneous parameters until the elastic regime ends and plastic growth occurs. Then we divide by two the elastic modulus of a small group of cells (marked with a white star in Fig. 3a) that will be referred to as “bump cells” thereafter. All the details of the computations are presented in SI. First, Fig. 3b shows that the multicellular system grows globally in the same way as the single hexagonal cell studied above; it diverges from the Lockhart predictions because the ratio *A/V* of the cells is not constant: the (ALPHA+) simulations exhibit a very large initial growth rate that decreases only when the cells are so large that water fluxes become limiting. The (PM-) set leads to a roughly twice lower growth rate than (REF). The set (CC-) leads to the same dynamics at the tissue level as (REF), because the total influx of water is not affected by fluxes between cells in this setup.

**Fig. 3.**
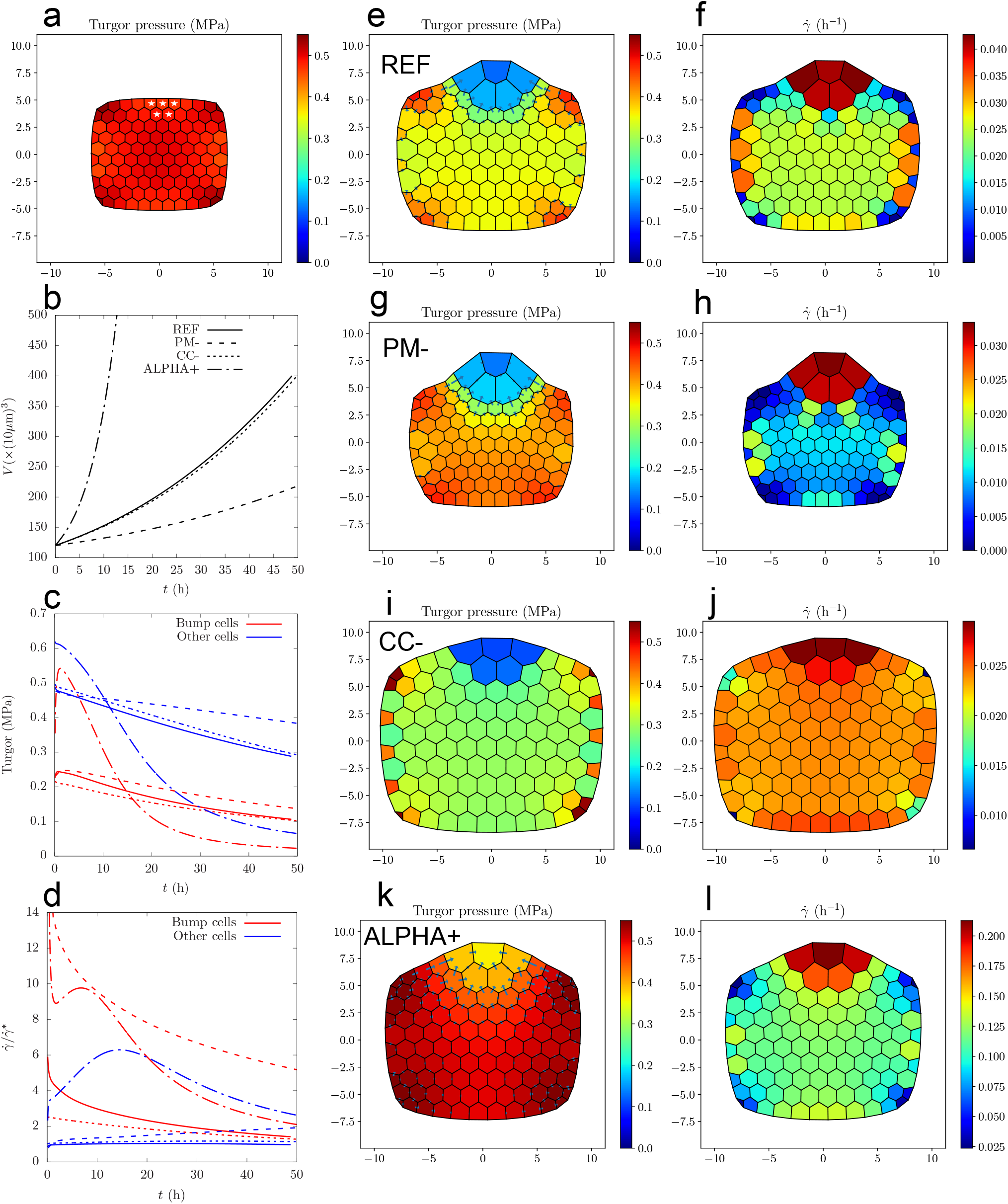
Growth of tissue with heterogeneous mechanical parameters, see table 1 in SI. a) **(a)** Initial state for (REF): walls are under tension because of turgor and have reached their yield deformation. At *t* = 0, the walls of the cells marked with a white star are softened (the elastic modulus is divided by two). **(b)** Time evolution of the total volume. The dashtype of the lines distinguishes the parameters sets; the same dashtype convention is used in **(c)** and **(d)**. **(c)** Time evolution of turgor pressure of bump cells (red) and other cells (blue). **(d)** Time evolution of relative growth rate of bump cells (red) and other cells (blue). **(e-l)** Turgor and relative growth rate maps of parameters sets (REF) (**(e-f)**), (PM-) (**(g-h)**), (CC-) (**(i-j)**), and (ALPHA+) (**(k-l)**), at the time when the volume of the bump cells has increased by a factor 5: *t* = 51h for (REF), *t* = 33h for (PM-), *t* = 80h for (CC-), *t* = 14.8h for (ALPHA+). The arrows represent the intensity and direction of cell-cell water fluxes; the scale for arrows is the same for (REF), (PM-) and (CC-) and close to 4 times higher for (ALPHA+).

Then we turn to the observation of heterogeneities: we focus on the differences between the bump region and the rest of the tissue. For all the parameters sets, Fig. 3c shows that turgor is in general lower in bump cells, but the gap varies depending on the parameters, as it has been predicted by the study of the two-cells model: compared to (REF), the heterogeneity in turgor is increased by a lower cell-cell conductivity (set CC-), and decreased by a larger value of *α^a^* (set ALPHA+). Decreasing the value of *P^M^* (set PM-) does not alter much the turgor heterogeneity compared to (REF). The maps of turgor (Figs. 3e,g,i,k) confirm visually these observations.

Fig. 3d shows the time evolution of 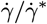 where 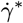 is the relative growth rate predicted by the Lockhart model (see Eq. (6)); its value is 2% h^-1^ for (REF), (CC-) and (ALPHA+), and 0.5% h^-1^ for (PM-). In the considered time frame, the relative growth rate of bump cells is always higher except for (ALPHA+): after an initial fast increase where bump cells grow faster, the tendency is inversed at *t* ≈ 20h because the bump cells have grown so much that fluxes become limiting. In the (REF) simulation, while the growth rate of non bump cells is almost constant and close to 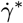, the growth rate of the bump cells is up to 6 times 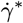 at the beginning of the simulation and progressively decreases toward 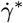. As a result of this large discrepancy, the bump region can be clearly distinguished from the rest of the tissue (Figs. 3e-f). In (CC-), the growth rate of the non bump cells is close to that of (REF), but the growth rate of the bump cells is much lower (Fig. 3d). As a result, the global shape remains convex and the bump is not clearly detached from the rest of the tissue (Figs. 3i-j). Note that (CC-) corresponds to a lower value of *α^s^* compared to (REF), which corresponded to a lower growth heterogeneity with the two-cells model studied above; this is also confirmed by the lower cell-cell fluxes towards the bump cells for (CC-), see the arrows in Figs. 3e,i. The (ALPHA+) simulation exhibits also a convex shape (Fig. 3k-l); it corresponds to a larger value of *α^a^* than (REF), and similarly to the two-cells model studied above, the growth rate heterogeneity is lower than (REF). Finally, the set (PM-) corresponds to an increase of the dimensionless parameter *ρ* (see Eq. (9)), and accordingly to an increase in growth rate heterogeneity as can be seen with Fig. 3d. Consequently, the bump region can clearly distinguished from the rest of the tissue, even better than (REF) (Fig. 3g-h); moreover, the growth of the cells close to the bump seems to be inhibited by fluxes as explained in the two-cells model described above and further explored below.

### Flux-based lateral inhibition predicted by the model

As we saw, cells that benefit from better mechanical conditions for growth (in the present case a lower elastic modulus) have a lower turgor than the other cells, and therefore attract water from them. Not only does it amplify their growth but it also inhibits the growth of their neighbours. Such a lateral inhibition mechanism is important for morphogenesis, as it allows very large growth rate heterogeneities and the appearance of well differentiated shapes (in the present case the appearance of a bump on the surface of the meristem). The efficiency of this mechanism varies depending on the position in the parameters space: for instance it is increased if the cell-cell conductivity *L^s^* (or equivalently *α^s^*) is increased (see Fig. 4a-d); even the whole tissue can be inhibited. Inhibited cells can also relax the tension of their walls and decrease their volume (see Fig. 4a). To further explore and quantify the spatial range of this inhibition process, we extended our two-cells model (see SI for detailed equations) to a chain of 2*N* + 1 cells where the central cell has twice softer walls. We numerically solved the corresponding system of differential equations for the set (REF) and then for a large range of values of *L^s^*. Fig. 4e shows that the number 2*N_i_* of inhibited cells scales with 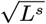. We computed the prefactor *c* (such that 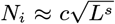) for values of (*α^a^*, *P^M^*) ∈ [0.05, 0.35] × [0.51, 0.85] (the interval for *P^M^* is in MPa) and plotted its value in the (*α^a^*, *P^M^*) space (Fig. 4f). This shows that the inhibition is favored by low values of *α^a^* and *P^M^* − *P^Y^*.

**Fig. 4.**
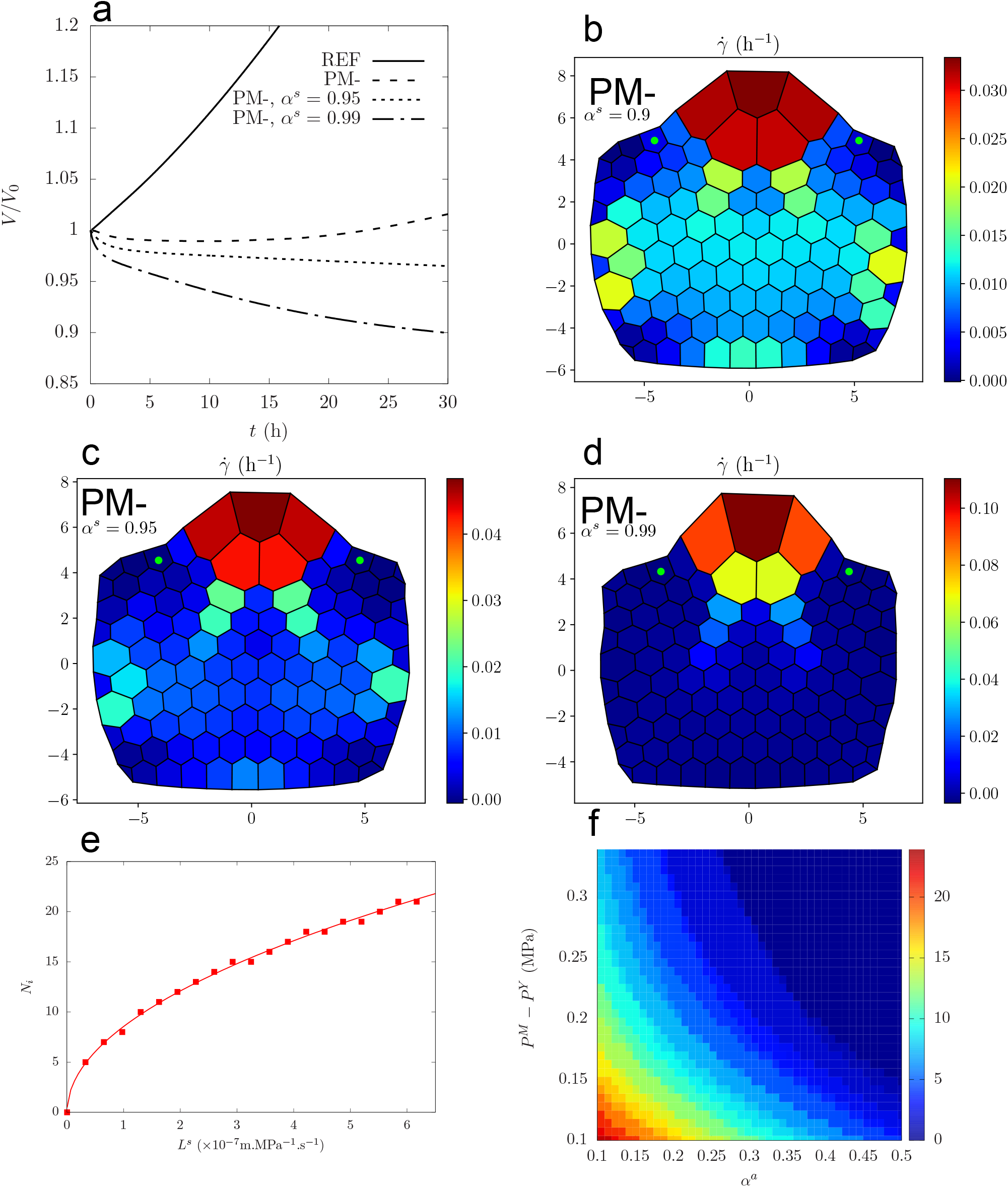
Evidence of lateral inhibition: left: **a)** time evolution of the volume of two cells on the boundary of the bump (marked with a green dot on the maps **b, c, d)** with the sets of parameters (REF), (PM-), (PM-) with *α^s^* = 0.95, (PM-) with *α^s^* = 0.99. *V*_0_ is the volume of the cells at *t* = 0. **b,c,d)** maps of relative growth rate at *t* = 33h for (PM-), *t* = 20h for (PM-) and α^s^= 0.95, *t*= 10h for (PM-) and *α^s^* = 0.99. **e-f)** Results for a chain of 2*N* + 1 cells with *N* = 50, where the central cell has twice softer walls; **e)** number *N_i_* of cells that are inhibited on each side of the central cell, for different values of *L^s^*; the line is a fit with a square root function, in the form 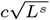. **f)** Values of the prefactor *c* in the space (*α^α^*, *P^M^*).

## Discussion

### A minimal model with a complex and rich behavior

The model proposed in this article is a minimal multicellular and multidimensional extension of the Lockhart 1-D single cell model; it can be regarded as a conceptual tool to study the interplay between fluxes and wall mechanics in a multicellular tissue. Wall expansion is modeled with a visco-elasto-plastic rheological law, while fluxes derive from water potential gradients. These two contributions are integrated into the mechanical equilibrium and interact through the pressure term. Contrary to most previous approaches, turgor is not an input of the model but a variable that adjusts simultaneously to mechanical, hydraulic, and geometrical constraints. First of all, this leads to a physically consistent representation of turgor: for instance, the model predicts that cells with softer walls have a lower turgor. Moreover, this has deep implications at tissue level: in the previous example, lower turgor is associated with a faster growth which can be itself amplified by fluxes that follow decreasing pressure gradients.

Thanks to the simplicity of the model, the predicted behavior can be analyzed and interpreted with two submodels built from the Lockhart model: first, a 1-D multicellular submodel was build with two or more side-by-side cells; it was used to study the growth of competing cells with heterogeneous properties. Key ingredients here are the wall synthesis threshold, the fact that fluxes and growth can relax turgor, and cell to cell fluxes that allow long range interactions. Second, in a 1-D system, cells are considered essentially as cylinders and their surface-to-volume ratio is constant. We thus extended also the Lockhart model in two dimensions, where cells have more degree of freedom to change their shape. In particular their allometric surface-to-volume ratio may then vary. This new possibility induces additional complexity in the tissue development as the rate of growth of cell surfaces may become a limiting factor for growing cells.

### A potentially new type of lateral inhibition mechanism

Depending on mechanical and hydraulic parameters of tissue regions, the model exhibits different growth regimes corresponding to either uniform or differential growth. One unexpected consequence of such an hydraulic-mechanical coupling at the tissue level is the observation that in certain regions of the parameter space where cell-to-cell hydraulic exchanges are non-limiting, growing tissue may exert an inhibiting influence on the growth of neighboring regions. This may be interpreted as a lateral inhibition mechanism. It has for long been recognized that lateral inhibitory mechanisms play a key role in setting some morphogenetic patterns in procaryotes (e.g. [28]), animals (e.g. [29, 30]) or plants (e.G. [31, 32]). Lateral inhibition operates in these systems via chemical signals, such as delta-notch in animals or auxin in plants. Our model predicts the existence of a novel type of lateral inhibition mechanism based on the coupling between mechanics and water fluxes. Previous observations of tissue growth suggest that such a phenomenon may occur in real tissues. In the shoot apical meristem for instance, detailed quantification of growth with cellular resolution indicates that the region surrounding primordia growth may have a negative growth rate [33], Figs. 2G and 3K. According to our model, this decrease of volume in boundary regions might be due to the primordium growth attracting locally most of the water supply and depriving lateral regions from water, and thus conforts the hypothesis of a new hydraulic-mechanical component of primordium lateral inhibition, beyond already identified auxin and cytokinin signals [34].

### Model simplifications and further potential extensions

Throughout the development of the model, we made several key choices concerning the abstraction of a multicellular plant tissue. First, our model was developed in 2-D for reasons of computational efficiency. In principle, it can be extended in 3-D, though at the expense of more complex formalism and implementation. Second, the current model considers that water transport is performed in the plant tissue through two conceptually different pathways ([1]). Water can first move within the apoplastic compartment between the cells and finally enter a cell. Water can also move locally from cell to cell. This movement includes itself conceptually both symplasmic movements (water circulates between cells through plasmodesmata without crossing membranes) and movements from cell to cell with intermediate steps in the wall (water is for example exported locally out of the cell by water transporters like aquaporins into the wall and immediately re-imported by water transporters into neighboring cells). For sake of simplicity in this first analysis, we represented the apoplasm as a single abstract compartment able to exchange water with every cell. To analyze precisely the effect of water transporters and their genetic regulation or to assess the impact of wall resistance to water movement in the processes, explicit spatial representation of the apoplasm, of plasmodesmata and of membrane water transporters could be integrated into the model in the future.

Finally, we considered a simplified situation here by imposing constant cell osmolarity. Allowing osmolarity variations (for instance higher values in faster growing regions) may impact turgor distribution (e.g [35]). However, this should not affect the ability of the system to build up growth heterogeneities. Similarly, we further simplified our model by keeping constant the apoplastic water potential. Relaxing this hypothesis would increase cell-cell water fluxes (via the apoplasm) and could also shift the model in the direction of the flux-limiting regime. This would therefore favor regimes where growth heterogeneities are amplified by fluxes.

## Supporting information

Supplementary info for the main article

## ACKNOWLEDGMENTS

This work has been carried out within the context of the project MecaFruit3D funded by the Agropolis foundation in Montpellier, France.

