## Supplementary info for the main article for "Coupling water fluxes with cell wall mechanics in a multicellular model of plant development"

#### This PDF file includes:

Figs. S1 to S3
Table S1
References for SI reference citations

DRAFT

### Supporting Information Text

#### 1. Calculations for simplified models

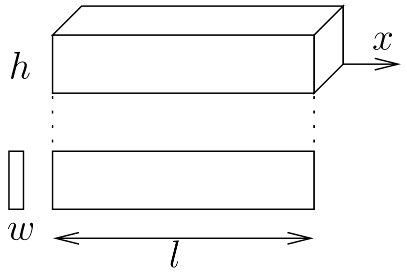


**Fig. S1.** Geometrical parameters of Lockhart-Ortega models: height $h$ and length $l$ of the cell, thickness $w$ of the walls. The two faces orthogonal to the $x$ axis are refered to as base faces while the four other faces are refered to as lateral faces.

**Lockhart-Ortega models.** The equations of cell wall elongation (Eq. (1) in main text) and of water uptake (Eq. 2 in main text) can be linked thanks to the geometry of the cell and the mechanical equilibrium. See Fig. S1 for the geometrical description.

First, the cell volume is $V = h^2 l$ and therefore we find that the relative growth rate of the cell is equal to the strain rate of the walls:

$$\dot{\gamma} = \frac{1}{V}\frac{\mathrm{d}V}{\mathrm{d}t} = \frac{1}{l}\frac{\mathrm{d}l}{\mathrm{d}t} = \dot{\varepsilon}. \qquad \text{[S1]}$$

Then, we consider the balance of forces on the base faces (see Fig. S1 for the nomenclature); their area is $h \times h$ and they are submitted to a total pressure force $Ph^2$ in the direction of the main axis of the cell, balanced by the tension from the lateral walls. Let $\sigma$ be the common (scalar) stress in the walls; the wall thickness is $w$ so their cross section is $h \times w$ and therefore they each exert a force $\sigma hw$ on the base faces. To be coherent with the bidimensional model we propose, we consider that the top and bottom lateral faces bear no stress and the balance of forces leads to

$$Ph^2 = 2\sigma hw$$

and therefore the balance of forces leads to $P = 2\frac{w}{h}\sigma$. Finally, thanks to this equation and the identity Eq. (S1), the Lockhart-Ortega model (eqs. (1), (3) in main text) is reduced to the following differential equation for $P$:

$$\frac{1}{\bar{E}}\frac{\mathrm{d}P}{\mathrm{d}t} + \phi^w (P - P^Y)_+ = \phi^a \left(P^M - P\right), \qquad \text{[S2]}$$

where $\phi^a = \frac{AL^a}{V}$ has been introduced in the main text; in order to keep the calculations as simple as possible, Lockhart made the assumption that the area of the base faces is negligeable compared to the area $A = 4hl$ of the lateral faces (see Fig. S1). Note that the cell volume is $V = h^2 l$ and therefore the ratio $A/V = 4/h$ is constant.

Let's study the transient behaviour of equation Eq. (S2), from an initial condition $P(t = 0) = 0$:

- Elastic regime: first, $P$ is below $P^Y$ and the plastic rate is zero; Eq. (S2) becomes

  $$\lambda^a \frac{\mathrm{d}P}{\mathrm{d}t} + P = P^M,$$

  where $\lambda^a = \frac{1}{\phi^a \bar{E}}$ is a characteristic time. The solution is

  $$P = P^M(1 - \exp(-t/\lambda^a)).$$

  The relative growth rate is

  $$\dot{\gamma} = \phi^a P^M \exp(-t/\lambda^a).$$

- Plastic regime: the plastic regime starts when $P = P^Y$, at $t^0 = \lambda^a \log\left(\frac{P^M}{P^M - P^Y}\right)$. The equation Eq. (S2) becomes:

  $$\frac{1}{\bar{E}}\frac{\mathrm{d}P}{\mathrm{d}t} + (\phi^a + \phi^w)P = \phi^a P^M + \phi^w P^Y,$$

and equivalently

$$\lambda^{aw}\frac{\mathrm{d}P}{\mathrm{d}t} + P = \alpha^a P^M + (1 - \alpha^a)P^Y,$$

DRAFT

where $\lambda^{aw} = \frac{1}{(\phi^a+\phi^w)\bar{E}}$ is a characteristic time. The solution is

$$P = \alpha^a P^M + (1-\alpha^a)P^Y - \alpha^a(P^M - P^Y)\exp((t^0 - t)/\lambda^{aw}), \tag{S3}$$

$$\dot{\gamma} = \frac{\phi^a\phi^w}{\phi^a+\phi^w}(P^M - P^Y) - \frac{(\phi^a)^2}{\phi^a+\phi^w}(P^M - P^Y)\exp((t^0 - t)/\lambda^{aw}). \tag{S4}$$

The stationnary solution is

$$P^* = \alpha^a P^M + (1-\alpha^a)P^Y \tag{S5}$$

$$\dot{\gamma}^* = \frac{\phi^a\phi^w}{\phi^a+\phi^w}(P^M - P^Y). \tag{S6}$$

**Single polygonal cell.**

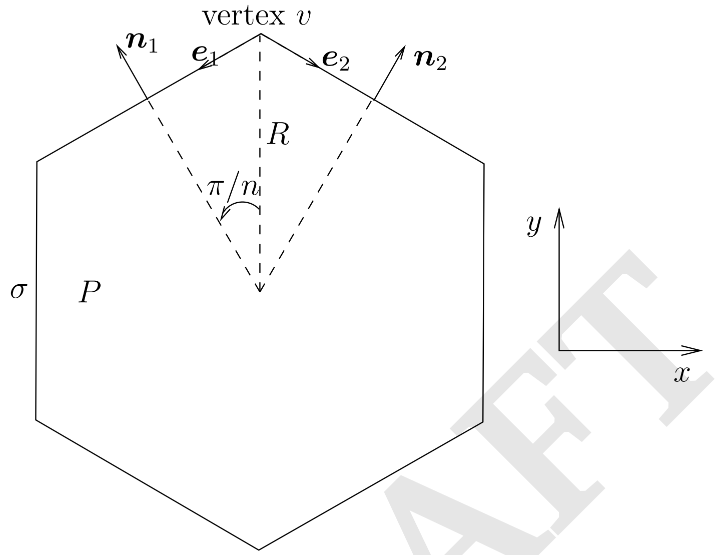


We consider a regular convex polygon of radius $R$ with $n$ edges that represents a cell.

**Mechanical equilibrium.** Let $\sigma$ be the stress in the walls and $P$ the pressure inside the cell; the outside pressure is set to zero. The length of the edges is $2R\sin(\pi/n)$, and the walls are given a height $h$ and a thickness $w$; therefore the stresses are exerted on a surface $hw$; the contribution of pressure on vertex $v$ is $\frac{1}{2}P2hR\sin(\pi/n)(\boldsymbol{n}_1+\boldsymbol{n}_2)$. Therefore, the balance of forces on vertex $v$ writes:

$$\frac{1}{2}P2hR\sin(\pi/n)(\boldsymbol{n}_1+\boldsymbol{n}_2) + \sigma hw(\boldsymbol{e}_1+\boldsymbol{e}_2) = 0.$$

The normal vectors are

$$\boldsymbol{n}_1 = (-\sin(\pi/n), \cos(\pi/n)) \quad \text{and} \quad \boldsymbol{n}_2 = (\sin(\pi/n), \cos(\pi/n)).$$

The tangent vectors are

$$\boldsymbol{e}_1 = (-\cos(\pi/n), -\sin(\pi/n)) \quad \text{and} \quad \boldsymbol{e}_2 = (\cos(\pi/n), -\sin(\pi/n)).$$

By symetry, the $x$ component of the resulting force is zero; the projection of the balance of forces on $y$ axis yields

$$2PhR\sin(\pi/n)\cos(\pi/n) - 2\sigma hw\sin(\pi/n) = 0,$$

and

$$P = \frac{w}{R\cos(\pi/n)}\sigma. \tag{S7}$$

When $n \to \infty$, $\cos(\pi/n) \to 1$ and we recover the Laplace law.

**Flux equation.** The surface of the polygon is

$$S_n = n \times 2R\sin(\pi/n)R\cos(\pi/n)/2 = R^2 n\sin(\pi/n)\cos(\pi/n).$$

The volume of the cell is $V = S_n h$, so the volume variation is

$$\frac{\mathrm{d}V}{\mathrm{d}t} = 2hR\frac{\mathrm{d}R}{\mathrm{d}t}n\sin(\pi/n)\cos(\pi/n).$$

DRAFT

The perimeter of the polygon is $n \times 2R\sin(\pi/n)$ so the lateral area of the cell is

$$A = 2nhR\sin(\pi/n).$$

Note that the ratio $A/V$ is not constant:

$$\frac{A}{V} = \frac{2}{R\cos(\pi/n)}.$$

Finally, the flux equation writes

$$2hR\frac{\mathrm{d}R}{\mathrm{d}t}n\sin(\pi/n)\cos(\pi/n) = n2hR\sin(\pi/n)L(P^M - P),$$

which yields

$$\frac{\mathrm{d}R}{\mathrm{d}t} = \frac{L}{\cos(\pi/n)}(P^M - P) \qquad \text{[S8]}$$

**Wall rheology.** Let $\varepsilon^e$ be the elastic deformation of the walls; it is related to the stress by the constitutive equation $\sigma = E\varepsilon^e$ where $E$ is the elastic modulus. The length of the edges is $l = 2R\sin(\pi/n)$ and therefore the strain rate of the edges is $\frac{1}{l}\frac{\mathrm{d}l}{\mathrm{d}t} = \frac{1}{R}\frac{\mathrm{d}R}{\mathrm{d}t}$. The rheological behaviour of the walls is given by

$$\frac{1}{R}\frac{\mathrm{d}R}{\mathrm{d}t} = \frac{\mathrm{d}\varepsilon^e}{\mathrm{d}t} + \Phi^w E\max(0, \varepsilon^e - \varepsilon^Y), \qquad \text{[S9]}$$

or equivalently

$$\frac{1}{R}\frac{\mathrm{d}R}{\mathrm{d}t} = \frac{1}{E}\frac{\mathrm{d}\sigma}{\mathrm{d}t} + \Phi^w\max(0, \sigma - \sigma^Y), \qquad \text{[S10]}$$

where $\varepsilon^Y$ (resp. $\sigma^Y$) is a yield elastic deformation (resp. stress).

**Numerical results.** The problem to solve is reduced to a set of two differential equations. It is numerically solved with the `odeint` routine from the `python` library `scipy`.

We study the growth of a hexagonal cell ($n = 6$) growing from an initial state where the elastic deformation of the walls is set to the threshold value, in order to bypass the pure elastic regime; computations are run over a long time scale. We want to study how this models compares to Lockhart-Ortega when the relative importance of fluxes and wall synthesis varies; to this end, we run three simulations with $\alpha^a = 0.1, 0.5, 0.9$. Let $R_0 = 10\mu$m be the initial radius of the cell, then $P^Y = \frac{w}{R_0\cos(\pi/6)}E\varepsilon^Y$ is a representative value for the yield turgor of a hexagonal cell. The value $\varepsilon^Y = 0.1$ is chosen accordingly to experimental observations where wall deformations can be of the order of 10%; then we choose $E$ such that $P^Y = 0.5$ MPa, which sets an order of magnitude for the initial turgor of the cell, close to observed experimental data. We choose $P^M = 0.7$ MPa so that it is above $P^Y$. Finally, we can use the Lockhart's prediction Eq. (S6) as an order of magnitude of the relative growth rate; we choose $\dot{\gamma}^* = 2\% \cdot \mathrm{h}^{-1}$. Then, a given value of $\alpha^a$ (evaluated with the initial area of the cell) sets a unique value of $L^a$ and $\phi^w$.

At the onset of the simulation, walls start to extend irreversibly and plastic growth occurs. Fig. S2a,c shows that the volume increases faster for large values of $\alpha^a$, although we have chosen the parameters so that the Lockhart model predicts a constant and common value of $\dot{\gamma}$. Fig. S2b shows that $P$ is initially close to Lockhart predictions $P^*$ but decreases fastly to zero; the fast decrease of $P$ coincides with peaks of $\dot{\gamma}$ (Fig. S2c) above the value $\dot{\gamma}^*$ with a higher peak for larger values of $\alpha^a$; the elastic deformation $\varepsilon^e$ (Fig. S2d)is not constant either, with a large peak above the Lockhart-Ortega prediction for $\alpha^a = 0.9$. For all values of $\alpha^a$, $\varepsilon^e$ converges toward the threshold $\varepsilon^Y$.

DRAFT

**Two-cells model.** The geometry and notations of the two-cells model is recalled in Fig. S3. Gathering the flux equation (Eq. 8 from main text) and the wall mechanics equation (Eq. 1 from main text) with $\frac{\mathrm{d}P}{\mathrm{d}t} = 0$, we get

$$\phi^a(P^M - P_0) + \frac{\phi^s}{2}(P_1 - P_0) - \phi^w(P_0 - P_0^Y)_+ = 0 \qquad \text{[S11]}$$

$$\phi^a(P^M - P_1) - \frac{\phi^s}{2}(P_1 - P_0) - \phi^w(P_1 - P_1^Y)_+ = 0. \qquad \text{[S12]}$$

First, we assume that both cells are growing ($P_i > P_i^Y, i = 0, 1$).

**First regime:** $P_i > P_i^Y, i = 0, 1$**.** Adding Eq. (S11) and Eq. (S12) we get:

$$\overline{P} = \alpha^a P^M + (1 - \alpha^a)\overline{P}^Y, \qquad \text{[S13]}$$

where $\alpha^a = \dfrac{\phi^a}{\phi^a + \phi^w}$, $\overline{P} = \dfrac{P_0 + P_1}{2}$. With Eq. 1 from main text, we get

$$\overline{\dot{\gamma}} = \frac{\phi^a\phi^w}{\phi^a + \phi^w}(P^M - \overline{P}^Y), \qquad \text{[S14]}$$

where $\overline{\dot{\gamma}} = \frac{\dot{\gamma}_0 + \dot{\gamma}_1}{2}$ Therefore, the gathering of two cells behaves the same as one cell if one considers the mean values.

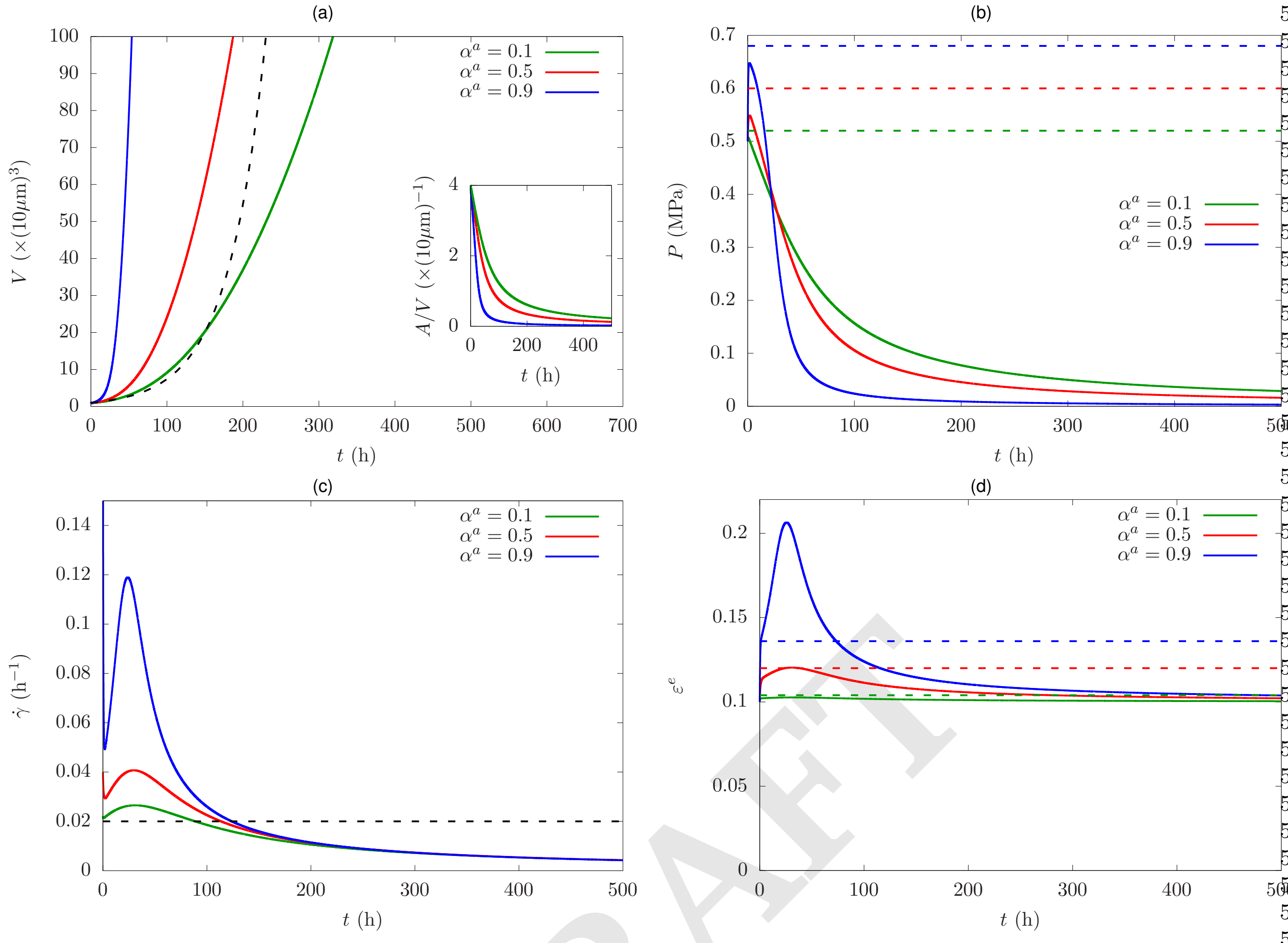


**Fig. S2.** Growth of a single hexagonal cell for three different values of $\alpha^a$: time evolution of volume (inset: ratio area/volume) (a), turgor (b), relative growth rate (c), and elastic deformation of the walls (d). The dashed lines correspond to the solution of the Lockhart model; note that the chosen sets of parameters lead to the constant and equal value $\dot\gamma^* = 2\% \cdot \mathrm{h}^{-1}$, and to the same evolution of volume.

DRAFT

Then, we examine the heterogeneities in turgor and growth rate. Substracting Eq. (S11) to Eq. (S12), we get

$$\Delta P = \frac{\phi^w}{\phi^a + \phi^s + \phi^w} \Delta P^Y.$$

Let

$$\alpha^s = \frac{\phi^s}{\phi^s + \phi^a}.$$

Then the previous expression becomes

$$\Delta P = \frac{(1-\alpha^a)(1-\alpha^s)}{1-\alpha^s+\alpha^a\alpha^s} \Delta P^Y. \quad \text{[S15]}$$

As $(1-\alpha^a)(1-\alpha^s) = 1 - \alpha^a - \alpha^s + \alpha^a\alpha^s < 1 - \alpha^s + \alpha^a\alpha^s$, we find that turgor difference $\Delta P$ cannot exceed the value $\Delta P^Y$. When $\alpha^s = 0$ (symplasmic fluxes negligeable with respect to apoplasmic ones), then $\Delta P = (1-\alpha^a)\Delta P^Y$; when $\alpha^s > 0$, symplasmic fluxes tend to reduce the turgor heterogeneity between cells.

With Eq. 7 from main text we get then

$$\Delta\dot\gamma = \frac{(\phi^a + \phi^s)\phi^w}{\phi^a + \phi^s + \phi^w} \Delta P^Y, \quad \text{[S16]}$$

where $\Delta\dot\gamma = \frac{\dot\gamma_0 - \dot\gamma_1}{2}$. Note that this expression is valid iff $P_1 > P_1^Y$ or equivalently $\dot\gamma_1 > 0$. The limit $\dot\gamma_1 = 0$ corresponds to the situation where cell 0 is growing in such a way that it prevents cell 1 to grow because of the symplasmic fluxes between them. We examine how this situation can occur depending on the values of the sumplasmic conductivity $\phi^s$ and the other parameters.

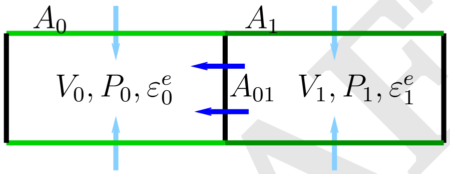


**Fig. S3.** Two cells model: symplasmic flows (dark blue arrows) occur through the contact surface $A_{01}$; apoplasmic flows (light blue arrows) occur through the surfaces $A_0$ and $A_1$. Growth is restricted to the green edges: cell $0$ (in dark green) has stiffer walls that cell 1 (in light green).

DRAFT

We find that

$$P_1 > P_1^Y \iff \frac{\phi^a + \phi^s}{\phi^a + \phi^s + \phi^w} \frac{\Delta P^Y}{P^M - \overline{P}^Y} < \frac{\phi^a}{\phi^a + \phi^w}$$
$$\iff \frac{\alpha^a}{1 - (1 - \alpha^s)\alpha^a} \rho < \alpha^a$$
$$\iff \alpha^s < \frac{1 - \rho}{1 - \alpha^a}.$$

For instance, $P_0^Y = 0.25$ MPa, $P_1^Y = 0.5$ MPa, and $P^M = 0.625$ MPa yields $\rho = 0.5$. The hypothesis of this study ($P_0^Y < P_1^Y < P^M$) corresponds to the condition $\rho \in [0, 1]$. Note that if $\alpha^a > \rho$, then $\frac{1-\rho}{1-\alpha^a} > 1$, and the condition is verified whatever the value of $\alpha^s$; if $\alpha^s = 1 - \rho$, the condition is equivalent to $\alpha^a > 0$, which is also always verified. Fig. S3a) recapitulates the regions of the parameters space $\alpha^a \times \alpha_s$ where the condition is verified, for different values of $\rho$. The size of the region $\dot{\gamma}_1 = 0$ increases as $\rho$ gets closer to 1.

**Second regime: $P_0 > P_0^Y$ and $P_1 < P_1^Y$.** In this case, eqs. Eq. (S11) and Eq. (S12) turn into

$$\phi^a(P^M - P_0) + \frac{\phi^s}{2}(P_1 - P_0) - \phi^w(P_0 - P_0^Y) = 0 \quad \text{[S17]}$$

$$\phi^a(P^M - P_1) - \frac{\phi^s}{2}(P_1 - P_0) = 0. \quad \text{[S18]}$$

Eq. (S18) leads to

$$P_1 = (1 - \tilde{\alpha}^s)P^M + \tilde{\alpha}^s P_0, \quad \text{[S19]}$$

where $\tilde{\alpha}^s = \frac{\phi^s}{2\phi^a + \phi^s}$. Adding eqs. Eq. (S17) and Eq. (S18) leads to

$$P_0(\phi^a + \phi^w) = 2\phi^a P^M + \phi^w P_0^Y - \phi^a((1 - \tilde{\alpha}^s)P^M + \tilde{\alpha}^s P_0),$$

then,

$$P_0(\phi^a(1 + \tilde{\alpha}^s) + \phi^w) = \phi^a(1 + \tilde{\alpha}^s)P^M + \phi^w P_0^Y,$$

and finally

$$P_0 = \alpha^{as} P^M + (1 - \alpha^{as})P_0^Y, \quad \text{[S20]}$$

where

$$\alpha^{as} = \frac{\phi^{as}}{\phi^{as} + \phi^w} \quad \text{and} \quad \phi^{as} = \phi^a(1 + \tilde{\alpha}^s).$$

Hence, thanks to the symplasmic fluxes from its neighbour cell 1, cell 0 benefits from an enhanced access to the apoplasmic fluxes by a factor $\phi^{as}/\phi^a = 1 + \tilde{\alpha}^s$. Then, from Eq. 1 in main text, the relative growth rate of cell 0 is

$$\dot{\gamma}_0 = \frac{\phi^{as}\phi^w}{\phi^{as} + \phi^w}(P^M - P_0^Y). \quad \text{[S21]}$$

By hypothesis, the growth rate of cell 1 is zero, and we can compute the heterogeneity in turgor: from Eq. (S19), we find that

$$\Delta P = \frac{1 - \tilde{\alpha}^s}{2}(P^M - P_0),$$

and hence

$$\Delta P = \frac{1}{2}(1 - \tilde{\alpha}^s)(1 - \alpha^{as})(P^M - P_0^Y). \quad \text{[S22]}$$

### 2. Numerical resolution of the 2D multicellular model

**Structure of the mathematical problem.** Thanks to the geometrical constraint of uni-directional growth, the Lockhart-Ortega is very simple to resolve. The identity between the relative growth rate of the cell and the strain rate of the walls allows to couple the equation that describes fluxes, and the equation that describes walls synthesis. Then the stress in the walls and the pressure inside the cell are linked by the mechanical equilibrium. Finally there is only one independent variable (pressure for instance) and the model can be solved analytically.

Conversely, in the bidimensionnal model we propose, the properties of a given wall (elongation rate and elastic deformation) cannot be directly linked to the properties of the adjacent cells (growth rate and pressure). Hence a new strategy has to be developped. First, we emphasize the strong coupling between fluxes and mechanics: the motion of the vertices is prescribed by the mechanical equilibrium (Eq. 11 from main text) between pressure forces and elastic forces; meanwhile, a displacement of the vertices can cause a variation of volume of several cells, which has to be balanced by water fluxes (Eq. 10 from main text); water fluxes are limited by the finite permeability of the walls, which sets a constraint on possible variations of volume. Similarly, any variation in the length of the walls leads to a modification of their elastic deformation (Eq. 7 from main text).

DRAFT

Another way to understand this problem is to consider it as the minimization of mechanical energy (mechanical equilibrium Eq. 11 from main text) under two constraints on the position of the vertices, through the volumes of the cells (Eq. 10 from main text) and the lengths of the edges (Eq. 7 from main text). This kind of problem is often encountered in mechanics, *e.g* solid friction, contact mechanics, or incompressible fluid mechanics; a powerfull theoritical and practical tool to solve this is the method of lagrangian multipliers. For instance, in the context of incompressible fluid mechanics, the constraint of volume conservation is relaxed by pressure that acts as a lagragian multiplier. Physically, the pressure adjusts itself so that both the constraint and the mechanical equilibrium are satisfied. The model we propose exhibits the same structure, as pressure will adjust to both fluxes and mechanical constraints. However, the system here is discrete, and the flux equation (Eq. 10 in main text) is linear with respect to pressure, so it can be reduced to a linear system. We will take advantage of this for the resolution of the model.

### Resolution algorithm.

**Volumes and lengths as functions of the positions of the vertices.** First, we express volumes and lengths as functions of the positions of the vertices. Let $N_v$ be the number of vertices and $\boldsymbol{X} \in \mathbb{R}^{2N_v}$ the vector of the positions of all the vertices. The volume of a cell $i$ is $V_i = S_i h$ where $S_i$ is its surface. As cells are non intersecting polygons, their signed surface is given by the general formula

$$S_i = \frac{1}{2} \sum_{k=0}^{n_i - 1} (x_k y_{k+1} - x_{k+1} y_k), \qquad \text{[S23]}$$

where $n_i$ is the number of vertices of cell $i$, $(x_k, y_k)_{k=0,\ldots,n_i-1}$ are the coordinates of the vertices of the cell $i$ in counterclockwise order, and we set $(x_{n_i}, y_{n_i}) = (x_0, y_0)$. Let $N_c$ be the number of cells and $\boldsymbol{V} \in \mathbb{R}^{N_c}$ the vector of all the cells volumes; thanks to Eq. (S23), it can be expressed as a function of $\boldsymbol{X}$ and its gradient $\nabla_{\boldsymbol{X}} \boldsymbol{V}$ with respect to $\boldsymbol{X}$ can be computed. Then the time derivative of $\boldsymbol{V}$ expresses as

$$\frac{\mathrm{d}\boldsymbol{V}}{\mathrm{d}t} = \nabla_{\boldsymbol{X}} \boldsymbol{V} \frac{\mathrm{d}\boldsymbol{X}}{\mathrm{d}t}.$$

Note here that $\nabla_{\boldsymbol{X}} \boldsymbol{V}$ is a $N_c \times 2N_e$ matrix and $\frac{\mathrm{d}\boldsymbol{X}}{\mathrm{d}t}$ is a $2N_e$ vector, so their product is well defined and has the correct dimension.

Similarly, the length of a segment $k$ with two vertices $v_1 = (x_1, y_1)$ and $v_2 = (x_2, y_2)$ at its ends is

$$l_k = \sqrt{(x_1 - x_2)^2 + (y_1 - y_2)^2}. \qquad \text{[S24]}$$

Let $N_e$ be the number of edges and $\boldsymbol{l} \in \mathbb{R}^{N_e}$ the vector of all the edges lengths; thanks to Eq. (S24), it can be expressed as a function of $\boldsymbol{X}$ and its gradient $\nabla_{\boldsymbol{X}} \boldsymbol{l}$ with respect to $\boldsymbol{X}$ can be computed. Then the time derivative of $\boldsymbol{l}$ expresses as

$$\frac{\mathrm{d}\boldsymbol{l}}{\mathrm{d}t} = \nabla_{\boldsymbol{X}} \boldsymbol{l} \frac{\mathrm{d}\boldsymbol{X}}{\mathrm{d}t}.$$

**Time discretisation.** Time is discretized using a fixed time step $\Delta t$ and the time derivatives are approximated by the 1st order Euler scheme, for instance:

$$\frac{\mathrm{d}\boldsymbol{X}}{\mathrm{d}t}(t) \approx \frac{\boldsymbol{X}(t + \Delta t) - \boldsymbol{X}(t)}{\Delta t}.$$

Let $\boldsymbol{\varepsilon} \in \mathbb{R}^{N_e}$ be the vector of all the elastic deformations of the edges. Let $\boldsymbol{X}^0 = \boldsymbol{X}(0)$ and $\boldsymbol{\varepsilon}^0 = \boldsymbol{\varepsilon}(0)$ be some initial conditions. We construct successive approximations of the solution at times $t_n = n\Delta t$ for $n > 0$ by solving at each time step the mechanical equilibrium (Eq. 11 from main text) along with the discretized versions of flux (Eq. 10 from main text) and wall rheology (Eq. 7 from main text) equations: let $\boldsymbol{P} \in \mathbb{R}^{N_c}$ be the vector of all the cells pressures; these equations can be written in a matrix form:

$$\nabla_{\boldsymbol{X}} \boldsymbol{V}(\boldsymbol{X}^{n+1}) \frac{\boldsymbol{X}^{n+1} - \boldsymbol{X}^n}{\Delta t} = M_P \boldsymbol{P}^{n+1} + \boldsymbol{b}_P, \qquad \text{[S25]}$$

$$\frac{\boldsymbol{\varepsilon}^{n+1} - \boldsymbol{\varepsilon}^n}{\Delta t} + \boldsymbol{\beta}^n \boldsymbol{\varepsilon}^{n+1} = \frac{1}{\boldsymbol{l}(\boldsymbol{X}^{n+1})} \nabla_{\boldsymbol{X}} \boldsymbol{l}(\boldsymbol{X}^{n+1}) \frac{\boldsymbol{X}^{n+1} - \boldsymbol{X}^n}{\Delta t}. \qquad \text{[S26]}$$

where $M_P$ is a $N_i \times N_i$ matrix, with the following non-zero coefficients:

$$M_P(i,i) = A_i L_i^a - \sum_{j \in n(i)} A_{ij} L_{ij}^s, \qquad \forall i = 1, \ldots, N_c,$$

$$M_P(i,j) = A_{ij} L_{ij}^s, \quad \forall i = 1, \ldots, N_c, \qquad \forall j \in n(i),$$

with $\boldsymbol{b}_P \in \mathbb{R}^{N_c}$ is defined by its coefficients

$$\boldsymbol{b}_p(i) = A_i L_i^a P^M, \qquad \forall i = 1, \ldots, N_c.$$

Note here that the model implies no time derivative of the pressure, so that $\forall n > 0$, $\boldsymbol{P}^{n+1}$ can be computed without the knowledge of $\boldsymbol{P}^n$, and the initial value of the pressure is not needed.

In addition, $\beta^n$ is the $N_e \times N_e$ diagonal matrix with components $\boldsymbol{\beta}^n(k,k) = \frac{2w}{h} \phi_k^w E_k \max\left(0, \frac{\varepsilon_k^n - \varepsilon_k^Y}{\varepsilon_k^n}\right)$ for $k = 1, \ldots, N_e$, and for the purpose of notation, $\frac{1}{\boldsymbol{l}}$ is the $N_e \times N_e$ diagonal matrix with components $1/l_k$. Note here that the variables $\boldsymbol{\beta}^n$ are taken at time step $n$ so that they are considered as constants at time step $n+1$ and the equation Eq. (S26) is linear with respect to the unknown $\boldsymbol{\varepsilon}^{n+1}$.

DRAFT

**Pressure and elastic deformation as functions of the position of the vertices.** Thanks to this time discretization, we see that at each time step, the unknown pressure $\boldsymbol{P}^{n+1}$ and elastic deformation $\boldsymbol{\varepsilon}^{n+1}$ are defined through the linear equations Eq. (S25) and Eq. (S26) which can be easily inverted, which allows to express both these variables as functions of the spatial unknown $\boldsymbol{X}^{n+1}$.

First, from equation Eq. (S25):

$$P(\boldsymbol{X}^{n+1}) = \frac{1}{\Delta t} M_P^{-1} \nabla_{\boldsymbol{X}} \boldsymbol{V}(\boldsymbol{X}^{n+1}) \boldsymbol{X}^{n+1} - M_P^{-1} \left( \frac{1}{\Delta t} \nabla_{\boldsymbol{X}} \boldsymbol{V}(\boldsymbol{X}^{n+1}) \boldsymbol{X}^{n} - \boldsymbol{b}_P \right). \quad \text{[S27]}$$

Then, using Eq. (S26):

$$\boldsymbol{\varepsilon}\left(\boldsymbol{X}^{n+1}\right) = \frac{1}{\Delta t} M_\varepsilon^{-1} \frac{1}{\boldsymbol{l}(\boldsymbol{X}^{n+1})} \nabla_{\boldsymbol{X}} \boldsymbol{l}(\boldsymbol{X}^{n+1}) \boldsymbol{X}^{n+1} - \frac{1}{\Delta t} M_\varepsilon^{-1} \left( \frac{1}{\boldsymbol{l}(\boldsymbol{X}^{n+1})} \nabla_{\boldsymbol{X}} \boldsymbol{l}(\boldsymbol{X}^{n+1}) \boldsymbol{X}^{n} - \boldsymbol{\varepsilon}^{n} \right), \quad \text{[S28]}$$

where $M_\varepsilon = \frac{1}{\Delta t} I_{N_e} + \boldsymbol{\beta}^n$.

**Structure of the resolution algorithm** Thanks to the two previous steps, we are now able to propose a algorithm for the resolution of the model.

- Initialization: Define $\boldsymbol{X}^0 \in \mathbb{R}^{2N_v}$ and $\boldsymbol{\varepsilon}^0 \in \mathbb{R}^{N_e}$
- $\forall n \geq 0$, assuming $\boldsymbol{X}^n$ and $\boldsymbol{\varepsilon}^n$ are known, let $\boldsymbol{F}^n : \mathbb{R}^{2N_v} \to \mathbb{R}^{2N_v}$ be the function such that $\forall v = 0, \dots, N_v - 1$,

  $$\begin{pmatrix} F^n_{2v+1}(\boldsymbol{X}) \\ F^n_{2v+2}(\boldsymbol{X}) \end{pmatrix} = \frac{1}{2} \sum_{k \in f(v)} \Delta_k P(\boldsymbol{X})\ A_k(\boldsymbol{X}) \boldsymbol{n}_k(\boldsymbol{X}) + \sum_{k \in f(v)} E_k \varepsilon^e_k(\boldsymbol{X}) a_k(\boldsymbol{X}) \boldsymbol{e}_{k,v}(\boldsymbol{X}),$$

  where $F^n_k$ is the $k$-th component of $\boldsymbol{F}^n$, and with the same notations as in Eq. 11 from main text; $\boldsymbol{P}(\boldsymbol{X})$ and $\boldsymbol{\varepsilon}(\boldsymbol{X})$ are the functions of $\boldsymbol{X}$ given by Eq. (S27) and Eq. (S28). Then, the new position of the vertices $\boldsymbol{X}^{n+1}$ is the solution of the equation

  $$\boldsymbol{F}^n(\boldsymbol{X}) = 0. \quad \text{[S29]}$$

**Resolution of** Eq. (S29)**.** This is the last and most critical step of the resolution algorithm. The problem of computing the roots of a multidimensional non linear function is often encountered in the mechanical modelling of complex multibody systems, and a method of choice for the resolution is the Newton algorithm [1]. It is a iterative process which derives from a Taylor expansion about a current point $\boldsymbol{u}^k$:

$$\boldsymbol{F}^n(\boldsymbol{u}^{k+1}) = \boldsymbol{F}^n(\boldsymbol{u}^k) + J(\boldsymbol{u}^k)(\boldsymbol{u}^{k+1} - \boldsymbol{u}^k) + o(\boldsymbol{u}^{k+1} - \boldsymbol{u}^k),$$

where $J(\boldsymbol{u}^k)$ is the jacobian matrix of function $\boldsymbol{F}^n$. The new value $\boldsymbol{u}^{k+1}$ is obtained by setting the right-hand side to zero and neglecting the high order term, and then solving the linear system:

$$J(\boldsymbol{u}^k)\delta\boldsymbol{u}^k = -F^n(\boldsymbol{u}^k), \boldsymbol{u}^{k+1} = \boldsymbol{u}^k + \delta\boldsymbol{u}^k.$$

With the initial value $\boldsymbol{u}^0 = \boldsymbol{X}^n$, iterations are run until a stopping criterium is met, for instance

$$\frac{\|\boldsymbol{F}^n(\boldsymbol{u}^k)\|}{\|\boldsymbol{F}^n(\boldsymbol{u}^0)\|} \leq tol_{res}, \quad \text{[S30]}$$

where $tol_{res} > 0$ is a fixed value. Then one can set $\boldsymbol{X}^{n+1} = \boldsymbol{u}^k$.

The compution of the jacobian matrix $J(\boldsymbol{u}^k)$ is non trivial here because of the numerous non-linearities of function $\boldsymbol{F}^n$. Therefore we have chosen to use the Newton-Krylov variant of this algorithm, that avoids the computation of the jacobian without loosing efficiency [1].

However, Newton methods in general have only local convergence properties, which means that they need an initial guess close enough to the solution to be able to converge. This is critical for instance in the first time step of the simulation, because the initial conditions might be far from equilibrium, but also for further time steps. This lack of global convergence properties is often dealt with by adding a friction term proportional to the velocity and hence to the time derivative of the positions. With this method, the problem to solve at each time step becomes after time discretization: find $\boldsymbol{X}$ such that

$$\boldsymbol{G}(\boldsymbol{X}) = \boldsymbol{F}^n(\boldsymbol{X}) - c\frac{\boldsymbol{X} - \boldsymbol{X}^n}{\Delta t} = 0,$$

where $c > 0$ is a friction coefficient. This new problem is easier to solve with the Newton method, all the more that $c$ is large. However, the root of $\boldsymbol{G}$ might not satisfy the condition Eq. (S30), and in addition its value depends on the value of $c$. Therefore, instead of applying the Newton method to the function $\boldsymbol{G}$, we perform the following iterative process:

- Initialization: $\boldsymbol{u}^0 = \boldsymbol{X}^n$

DRAFT

- Assuming $\boldsymbol{u}^k$ is known, compute $\boldsymbol{u}^{k+1}$ as the solution of

$$\boldsymbol{G}^k(\boldsymbol{u}^{k+1}) = 0, \tag{S31}$$

  where $\boldsymbol{G}^k(\boldsymbol{u}^{k+1}) = \boldsymbol{F}^n(\boldsymbol{u}^{k+1}) - c^k \frac{\boldsymbol{u}^{k+1} - \boldsymbol{u}^k}{\Delta t}$, and the value $c^k > 0$ will be adjusted to ensure a robust convergence (see below). This solution is computed thanks to the Newton method, with the tolerance $tol_{res}/10$ in the stopping criterium.

- The iterations are stopped when $\frac{\|\boldsymbol{F}^n(\boldsymbol{u}^k)\|}{\|\boldsymbol{F}^n(\boldsymbol{u}^0)\|} \leq tol_{res}$. Then the choice $\boldsymbol{X}^{n+1} = \boldsymbol{u}^k$ is an approximate solution of Eq. (S29).

In this algorithm, the choice of the friction coefficient $c^k$ is not straightforward: a large value would ensure the convergence of subproblem Eq. (S31), but it would also slow down the convergence toward the solution of problem Eq. (S29). To avoid this, we choose a large initial value $c^0$ and decrease it with the law $c^{k+1} = c^k/2$. This choice ensures a robust behaviour of the algorithm.

### 3. Sets of parameters used for the bump simulations

Let $R_0 = 10\mu$m be the initial radius of the cell, then $P^Y = \frac{w}{R_0 \cos(\pi/6)} E\varepsilon^Y$ is a representative value for the yield turgor of a hexagonal cell. However we have observed that the effective threshold pressure is approximately twice lower in multicellular tissues and we have adapted the value of $E$ accordingly: we choose $E$ such that $P^Y = 0.5$ MPa and multiplied this value by two to obtain a an order of magnitude for the initial turgor of the cell close to the target value 0.5 MPa. The value $\varepsilon^Y = 0.1$ is chosen accordingly to experimental observations where wall deformations can be of the order of 10%. We choose two values for $P^M$: 0.55 MPa close to the threshold, and 0.7 MPa. Finally, we can use the Lockhart's prediction $\dot{\gamma}^*$ (Eq.6 from main text) as an order of magnitude of the relative growth rate; we choose $\dot{\gamma}^* = 2\% \cdot \text{h}^{-1}$. Then, a given value of $\alpha^a$ (evaluated with $R = R_0$) sets a unique value of $L^a$ and $\phi^w$. The table S1 recapitulates the sets of parameters used in this article, either with the control parameters

$$\varepsilon^Y, P^M, P^Y, \dot{\gamma}^*, \alpha^a, \tag{S32}$$

or equivalently with the actual parameters of the model

$$\varepsilon^Y, P^M, E, \Phi^w, L^a. \tag{S33}$$

The correspondance has been obtained with $R_0 = 6.5\mu$m.

DRAFT

**Table S1. Parameters used for the bump simulation (see Fig. 3 in main text). The top part of the table refers to the control parameters** Eq. (S32)**, and the bottom part to the actual parameters** Eq. (S32) **used in the 2D model. The rightmost parameters after the vertical double bar are specific to multicellular models as they quantify the water conductivity between neighbour cells. The geometrical parameters are** $h = 10\mu\text{m}$ **and** $w = h/20$**.**

| **Control parameters** | $\varepsilon^Y$ | $P^M$ (MPa) | $P_6^Y$ (MPa) | $\dot{\gamma}^*$ ($\text{h}^{-1}$) | $\alpha^a$ | $\alpha^s$ |
|---|---|---|---|---|---|---|
| (REF) | 0.1 | 0.7 | 0.5 | $2 \cdot 10^{-2}$ | 0.1 | 0.9 |
| (CC-) | 0.1 | 0.7 | 0.5 | $2 \cdot 10^{-2}$ | 0.1 | 0.1 |
| (ALPHA+) | 0.1 | 0.7 | 0.5 | $2 \cdot 10^{-2}$ | 0.9 | 0.9 |
| (PM-) | 0.1 | 0.55 | 0.5 | $0.5 \cdot 10^{-2}$ | 0.1 | 0.9 |

| **Actual parameters** | $\varepsilon^Y$ | $P^M$ (MPa) | $E$ (MPa) | $\Phi^w$ ($\text{MPa}^{-1}.\text{s}^{-1}$) | $L^a$ ($\text{m.MPa}^{-1}.\text{s}^{-1}$) | $L^s$ ($\text{m.MPa}^{-1}.\text{s}^{-1}$) |
|---|---|---|---|---|---|---|
| (REF) | 0.1 | 0.7 | 112.6 | $2.8 \cdot 10^{-5}$ | $8.7 \cdot 10^{-11}$ | $7.8 \cdot 10^{-10}$ |
| (CC-) | 0.1 | 0.7 | 112.6 | $2.8 \cdot 10^{-5}$ | $8.7 \cdot 10^{-11}$ | $9.6 \cdot 10^{-12}$ |
| (ALPHA+) | 0.1 | 0.7 | 112.6 | $3.1 \cdot 10^{-6}$ | $7.8 \cdot 10^{-10}$ | $7.0 \cdot 10^{-9}$ |
| (PM-) | 0.1 | 0.55 | 112.6 | $2.8 \cdot 10^{-5}$ | $8.7 \cdot 10^{-11}$ | $7.8 \cdot 10^{-10}$ |

DRAFT
